# A community-level investigation of the yellow fever virus outbreak in South Omo Zone, South-West Ethiopia, 2012–2014

**DOI:** 10.1101/320317

**Authors:** Ranya Mulchandani, Fekadu Massebo, Fekadu Bocho, Claire L Jeffries, Thomas Walker, Louisa A Messenger

## Abstract

**Background:** A yellow fever (YF) outbreak occurred in South Omo Zone, Ethiopia in 2012-2014. This study aimed to analyse historical epidemiological data, to assess the risk for future YF outbreaks through entomological surveillance, including mosquito species identification and molecular screening for arboviruses, and finally to determine the knowledge, attitudes and current preventative practices within the affected communities.

**Methodology/Principal Findings:** From October 2012 to March 2014, 165 cases and 62 deaths were reported, principally in rural areas of South Ari region (83.6%), south-west Ethiopia. The majority of patients were 15-44 years old (74.5%) and most case deaths were males (76%). Between June and August 2017, 688 containers were sampled from across 177 households to identify key breeding sites for *Aedes* mosquitoes. *Ensete ventricosum* (“false banana”) was identified as the primary natural breeding site, and clay pots outside the home as the most productive artificial breeding site. Entomological risk indices from the majority of sites were classified as “high risk” for future outbreaks under current World Health Organization criteria. Adult trapping resulted in the identification of members of the *Aedes simpsoni* complex in and around households. Screening of adult females revealed no detection of yellow fever virus (YFV) or other arboviruses. 88% of 177 participants had heard of YF, however many participants easily confused transmission and symptoms of YF with malaria, which is also endemic in the area.

**Conclusions/Significance:** Study results emphasise the need for further entomological studies to improve our understanding of local vector species and transmission dynamics. Disease surveillance systems and in-country laboratory capacity also need to be strengthened to facilitate more rapid responses to future YF outbreaks.

**Author Summary:** Despite the availability of a highly effective vaccine, yellow fever virus (YFV) remains an important public health problem across Africa and South America due to its high case-fatality rate. This study aimed to assess and reduce the risk for future outbreaks. During this study, historical data analysis was conducted to understand the epidemiology of the recent outbreak in 2012-2014. Entomological surveillance was also carried out, including both mosquito species identification and molecular screening for arboviruses, as well as a household survey to understand the knowledge and attitudes towards yellow fever (YF) within the affected areas and to assess community-level practices for YF prevention. We found a high abundance of *Aedes simpsoni* complex in the context of low vaccination coverage. Community knowledge and practice levels were low for reducing potential breeding sites, highlighting the need for increased dissemination of information to community members on how to reduce their risk of exposure to mosquito vectors of arboviruses.

## Introduction

Yellow fever virus (YFV) is a flavivirus transmitted primarily to humans and non-human primates through the bite of an infected female *Aedes* spp. or *Haemagogus* spp. mosquito (1). YFV is endemic to Africa and Latin America where it causes a spectrum of clinical symptoms ranging in severity from asymptomatic infection, mild illness with flu-like symptoms to severe disease including, fever, jaundice or haemorrhage and death (2). Despite the availability of a highly effective vaccine, YFV continues to occur in epidemic situations, and it is estimated to result in 130,000 human cases and 78,000 deaths annually in Africa alone (2). Since the 1980s, there has been an unprecedented rise in the number of large YFV outbreaks (3), including Angola and the Democratic Republic of Congo in 2016 which together became one of the largest outbreaks in Africa for more than 20 years (4–6). In early 2018, numerous cases were confirmed in Nigeria and Brazil, which was particularly concerning as they were being reported from areas previously not considered at-risk (7). YFV has also re-emerged across East Africa with outbreaks in Ethiopia in 2012 and Uganda in 2016 (8). Due to this global resurgence, YFV continues to be considered a significant threat to public health (9).

The resurgence of epidemic yellow fever (YF) and the increased risk of urban outbreaks is multi-factorial (10,11). Rapid urbanisation, population migration, climatic changes and increased travel have all been implicated in expanding the geographical range of YFV and driving mosquito vectors closer to human dwellings where unvaccinated individuals are often living in highly populated areas (12). A multi-faceted approach is necessary for YFV control, which includes strong laboratory and surveillance systems with rapid case reporting, appropriate case management, vector control and reactive and preventive vaccination campaigns (13).

Ethiopia has experienced numerous YFV outbreaks since the 1940s. Between 1960-1962, the largest YFV outbreak ever recorded in Africa occurred along the River Omo, Ethiopia (Gamo Gofa, Jinka and Kaffa regions) which resulted in approximately 200,000 human cases and 30,000 deaths. In 1966, YFV appeared in Arba Minch, in an area previously unaffected in the 1960 epidemic and therefore excluded from the mass vaccination campaign at the time. During this outbreak, 450 deaths were reported (2,200 human cases) and the outbreak was confirmed through serological and entomological testing (14). After almost a 50-year absence, YFV re-emerged in 2012. A reactive vaccination campaign commenced in June, 2013, which reached approximately 550,000 people across the at-risk population. There have been no further vaccination campaigns since this outbreak, and Ethiopia is one of the few remaining endemic countries that have not introduced the YF vaccine into their childhood immunization programme. Therefore, the country is classified as a top priority through the Eliminating Yellow Fever Epidemics (EYE) strategy, a coalition of partners led by the World Health Organization (WHO), UNICEF and Gavi, the Vaccine Alliance (15).

Knowledge, Attitudes and Practices (KAP) studies have been widely used to understand the community context of disease transmission, to help inform appropriate control and risk communication activities, with an overall aim to reduce barriers to the prevention of infectious diseases. In Ethiopia, there is a considerable paucity of such data on YFV, nor had a YFV outbreak occurred in the region for almost 50 years. To guide appropriate, prospective disease control interventions, this study aimed to collect historical epidemiological information of the 2012 YFV outbreak, to assess the risk for future outbreaks through entomological surveillance, including mosquito species identification and molecular screening for arboviruses, and finally to determine knowledge and attitudes towards YF within the affected communities following this outbreak and assess current community-level practices for YF prevention.

## Methods

### Study location

The study was conducted in South Omo Zone (SOZ), Ethiopia, which is located in south-west Ethiopia (Southern Nations Nationalities and People’s Region – SNNPR) across 5 selected kebeles (villages), between June and August 2017 (Figure 1). Aykamer, Shepe, Arkisha (South Ari woreda (region)), Hana (Salamago woreda) and Besheda (Hammer woreda) kebeles were selected as representative sites which reported varying numbers of cases during the 2012-2014 outbreak and were all targeted for vaccination during the emergency reactive campaign in 2013.

**Figure 1:**
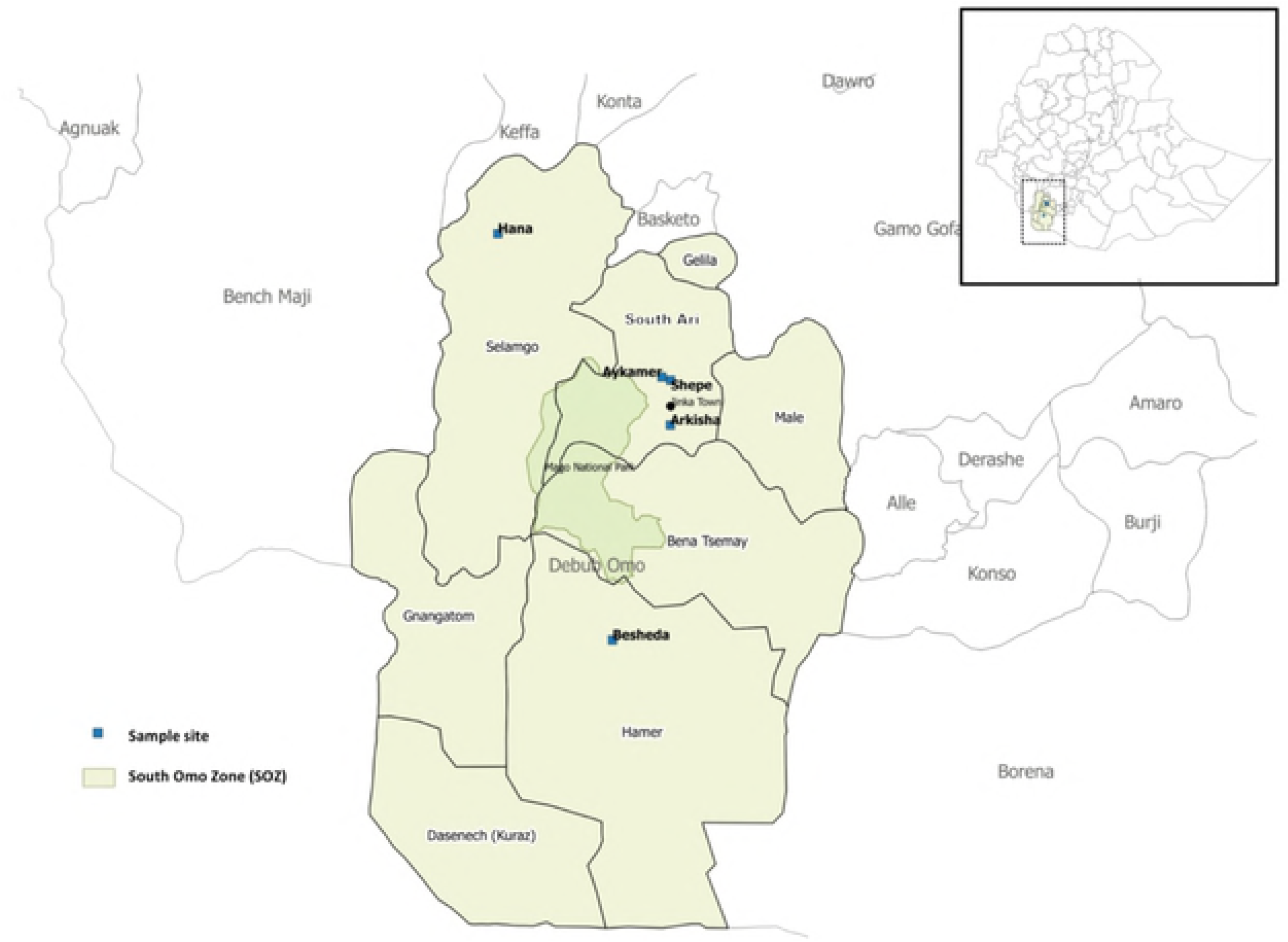
Map of study sites in South Omo Zone (SOZ), Southern Nations Nationalities and People’s Region (SNNPR), south-west Ethiopia.

### Historical clinical data collection

In November 2012, cases of an unknown febrile illness were reported to the Public Health Emergency Management (PHEM) through the mandatory weekly reporting format for health extension workers (HEW) on all immediately reportable diseases. Symptoms were a combination of fever, headache, nausea, bloody vomiting, abdominal pain, joint pain and jaundice. A team from the SOZ Health Department, WHO Ethiopia Country Office and Ethiopian Public Health Institute (EPHI) were deployed to the field for a rapid risk assessment. The causative agent was confirmed as YFV, following serum sample analysis. WHO standard case definition was used during the field investigation, and further cases confirmed through epi-link. The line list of all suspected and confirmed cases during the outbreak was retrieved from the SOZ Health Department in June 2017, to conduct a full historical descriptive epidemiological analysis; an interim report detailing cases up to October 2013 was published in 2017 by Lilay *et al*. (16).

### Entomological investigation and sampling

At each household premise, selected to participate in the KAP survey, immature *Aedes* spp. were sampled from all natural and artificial containers found around the home using a larval dipper. Only containers that contained water were included. An entomological survey was completed to define the type of container, its location, its usage and larval and pupal densities to calculate risk indices and to assess the most productive breeding sites. Any container that was found harbouring at least one larva or pupa of *Aedes* spp. was considered positive, and a sample of immature stages collected were reared to adulthood for subsequent morphological identification (17,18). The following risk indices were calculated:

*House Index (HI)* – the percentage of houses found positive for mosquito larvae or pupae. *Container Index (CI)* – the percentage of containers found positive for mosquito larvae or pupae. *Breteau Index (BI)* – the number of containers found positive for mosquito larvae and pupae per 100 houses surveyed. *Pupal Demographic Index (PDI)* – number of pupae found per number of residents in the houses inspected.

Entomological indices were interpreted according to the WHO guidelines; a high risk of YFV transmission is HI>35%, BI>50 or and CI>20% and low risk is HI<4%, CI<3% or BI<5 (19). The most productive containers were considered the types of water-holding containers where >70% of all pupae were found (20).

A Prokopack aspirator was used to collect adult mosquitoes at each household, both inside and outside of the premises. Each location was sampled for a total of 15 minutes (50% inside, 50% outside) to ensure systematic sampling. Specimens were labelled with time, date and location of their collection and stored in RNAlater® (Sigma, UK) at 4°C or lower to prevent viral RNA degradation.

### Molecular species identification

RNA was extracted from individual whole mosquitoes using QIAGEN RNeasy 96 kits according to manufacturer’s instructions. RNA was eluted in 40 μL of RNase-free water and stored at −80°C. A QIAGEN QuantiTect Reverse Transcription kit was used to reverse transcribe RNA to generate cDNA from all RNA extracts according to the manufacturer’s instructions. To determine the species of adult female *Aedes* collected in Shepe, Aykamer and Arkisha, a fragment of the ITS2 gene was sequenced (21). PCR products were separated and visualised using 2% E-Gel EX agarose gels (Invitrogen) with SYBR safe and an Invitrogen E-Gel iBase Real-Time Transilluminator. PCR products were submitted to Source BioScience (Source BioScience Plc, Nottingham, UK) for PCR reaction clean-up, followed by Sanger sequencing to generate both forward and reverse reads. Sequencing analysis was carried out in MEGA7 (22) as follows. Both chromatograms (forward and reverse traces) from each sample was manually checked, analysed, and edited as required, followed by alignment by ClustalW and checking to produce consensus sequences. Consensus sequences were used to perform nucleotide BLAST (NCBI) database queries and an alignment constructed to include all field sample consensus sequences; two consensus sequences from *Ae. bromeliae* specimens, generated by following the same procedure as the field samples; plus relevant sequences covering the region sequenced, obtained from GenBank. This alignment by ClustalW was used to produce a Maximum Likelihood (ML) phylogenetic tree based on the Tamura-Nei method (23).

### Arbovirus screening

Screening for YFV and other major arboviruses of public health suspected or having the potential of being transmitted in the region (dengue virus (DENV), Zika virus (ZIKV), chikungunya virus (CHIKV), West Nile virus (WNV) and Rift Valley fever virus (RVFV)) was undertaken using published real time PCR assays (Supplementary File S1). PCR reactions for all assays except ZIKV and WNV were prepared using 5 µL of QIAGEN SYBR Green Master mix, a final concentration of 1 µM of each primer, 1 µL of PCR grade water and 2 µL template cDNA, to a final reaction volume of 10 µL. Prepared reactions were run on a Roche LightCycler® 96 System and PCR cycling conditions are described in Supplementary File S1. Amplification was followed by a dissociation curve (95°C for 10 seconds, 65°C for 60 seconds and 97°C for 1 second) to ensure the correct target sequence was being amplified. ZIKV and WNV screening was undertaken using a Taqman probe-based assay using 5 µL of QIAGEN QuantiTect probe master mix, a final concentration of 1 µM of each primer, 1 µL of PCR grade water and 2 µL template cDNA, to a final reaction volume of 10 µL. PCR results were analysed using the LightCycler® 96 software (Roche Diagnostics). Synthetic long oligonucleotide standards (Integrated DNA technologies) of the PCR product template sequence were generated in the absence of biological virus cDNA positive controls and each assay included negative (no template) controls.

### Knowledge, Attitudes and Practices (KAP) study design and sample population

180 KAP surveys were administered across the five sites during the study period. A full household list was gathered from each village health post and random sampling used to select households, using a random number generator. Sample size calculations were conducted in STATA/IC 14.2 using the *svysampsi* command for surveys with a dichotomous outcome variable, with a proportion of 0.9 assumed to have the expected outcome, an error rate of 5%, a response rate of 90%, and a 95% confidence interval.

The KAP survey was developed through adaptation of previous KAP surveys used for other arboviruses (DENV and ZIKV), as well as compilation of context specific questions following informal discussions with stakeholders in Ethiopia (24–27). The survey was semi-structured, including both open and closed-ended questions, and captured details on household characteristics (socio-economic status/education), past YFV infection, vaccination status and general *Aedes* spp. control practices (Supplementary File S2). The survey was structured into eight main sections: [1] socio-demographic characteristics of study participants; [2] knowledge of YF symptoms, signs and transmission modes; [3] attitudes towards YF; [4] preventative practices against YF; [5] sources of information regarding YF; [6] YF case finding; [7] YF vaccination coverage estimation; and [8] other epidemiological risk factor information.

The full questionnaire was first developed in English, translated into Amharic and re-translated back into English for analysis. Before its use in the study, the questionnaire was pilot tested among community members in Arkisha, who were not included in the final analysis.

### KAP data collection and analysis

At each study site, the questionnaire was conducted in Amharic (or local dialect if preferred by respondent) by a HEW, who was trained by an experienced interviewer from Jinka Town. None of the interviewers were told the correct results to avoid interview bias during data collection. The head of household was the main respondent, however when this was not possible, another member of the family was interviewed *in lieu*.

All completed household surveys were double-checked and verified on the same day for completeness and consistency. Data were interpreted taking into account informal conversations and observations in the community and hospital.

The KAP assessment was conducted using a scoring system. A participant’s KAP score was calculated as the sum of their correct answers, where a correct answer was given a value of 1 and an incorrect answer (including any answers “do not know” or “no answer”) the value of 0 (25). The total possible score was 15 for Knowledge, 5 for Attitude and 8 for Practices. Respondents’ levels were defined as “good” or “poor” based on a 75% cut-off threshold. Logistic regression was conducted in STATA/IC 14.2 to identify determinants for KAP levels. Independent variables included in the model were household location, YF vaccination status, education level and the age and sex of respondent.

### Ethical approval

Ethical approval for the study was obtained from the London School of Hygiene and Tropical Medicine (LSHTM; ref #12291) and Arba Minch University (AMU; ref #6008/111). An invitation letter from AMU was taken to SOZ Health Department, who then provided an invitation letter, which was delivered to the village health posts to gain permission to sample. All participants were adults and gave full written informed consent. Confidentiality of all respondents was assured by using unique study identifiers. Working closely with HEWs ensured community acceptance and cooperation.

## Results

### Epidemiological information

Between November 2012 and March 2014, a total of 165 cases of YF were reported to PHEM, including 62 deaths and 4 laboratory confirmed cases (with the other 161 suspected cases defined through epi-link), in an entirely unvaccinated population. The index case was reported from Geza and first developed symptoms between 12^th^ and 23^rd^ November, 2012. Laboratory confirmation was on 7^th^ May, 2013. The main peak of the outbreak occurred from March to May, 2013, with the cases appearing to decline following the emergency vaccination campaign which commenced on 10^th^ June, 2013 (Figure 2).

**Figure 2:**
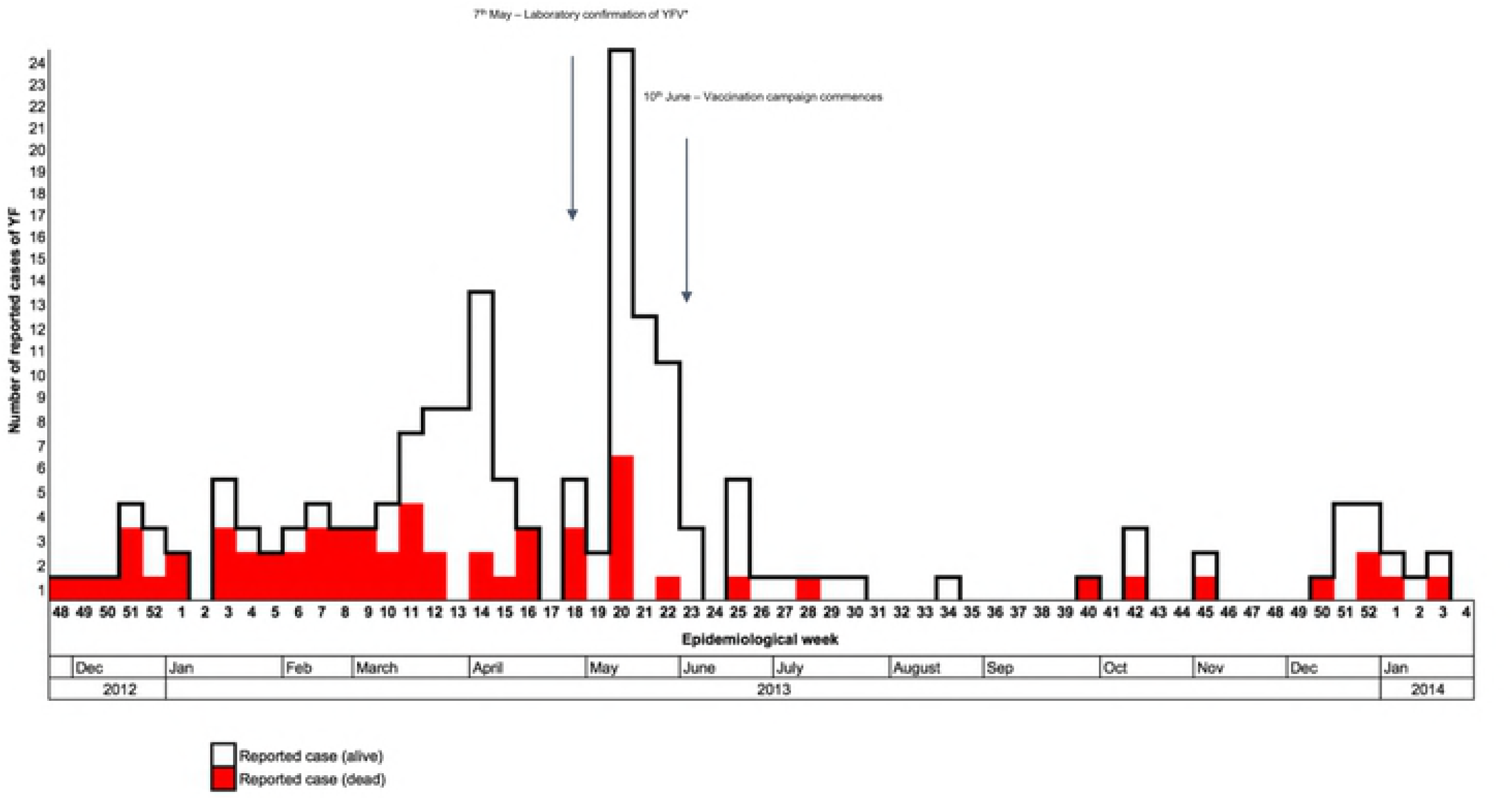
Distribution of reported yellow fever cases by their date of onset, from November 2012 to January 2014 (n=165) in Southern Nations, Nationalities, and People’s Region (SNNPR), Ethiopia. *YFV: Yellow fever virus.

The majority of cases were 15-44 year olds (75.8%), with males slightly more affected than females, with a male to female case ratio of 1.4:1 (Table 1). Most case deaths were also in males (76%), which resulted in a case fatality rate (CFR) of 48.5% compared to 22.1% in females (Table 2). The overall CFR was 37.6%, with the majority of deaths occurring at the start of the outbreak. 69% of reported deaths occurred in health facilities; the remaining 31% were community deaths found through active case detection.

**Table 1:**
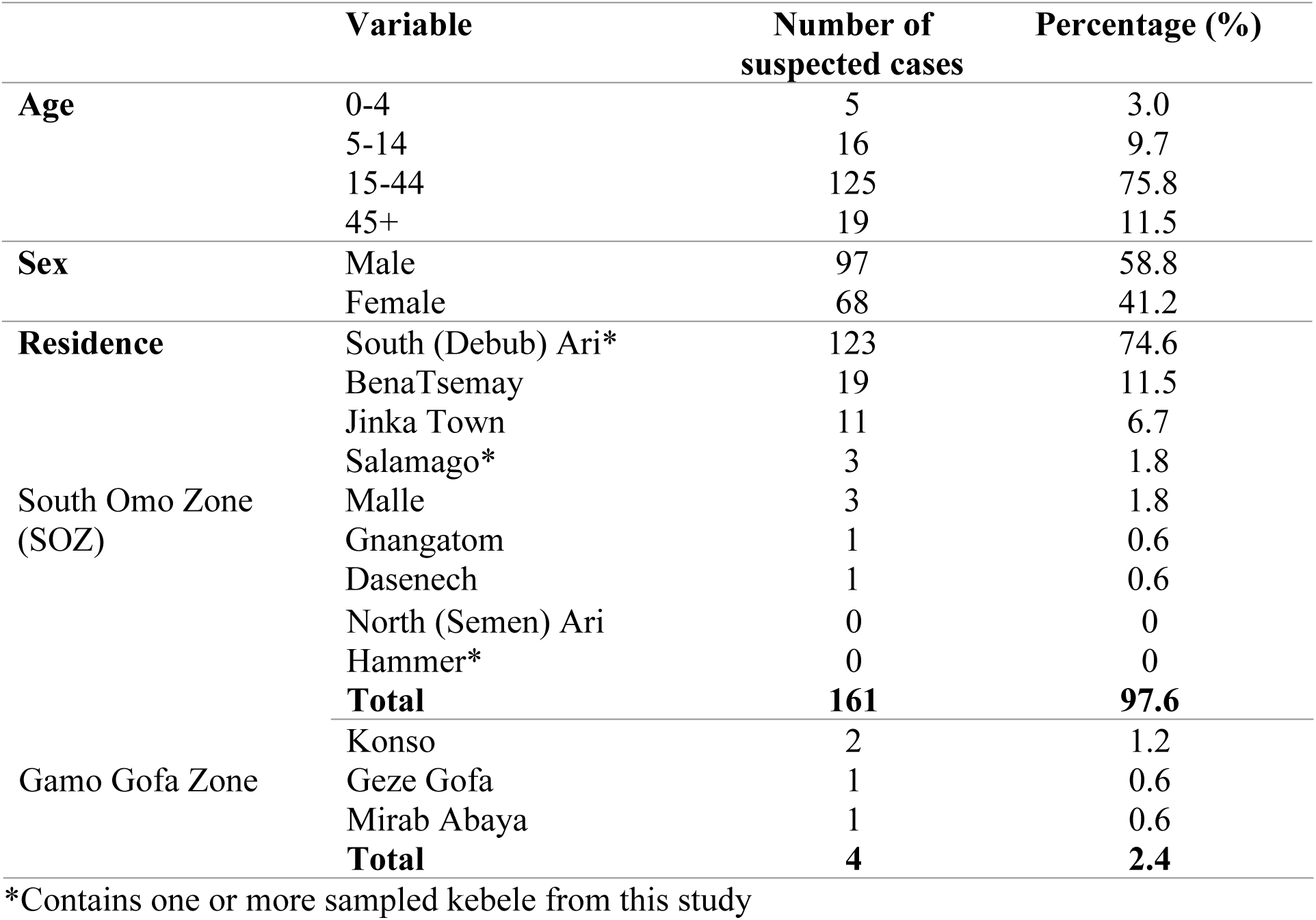
Age, sex and residence of reported yellow fever (YF) cases during the YF outbreak in Southern Nations, Nationalities, and People’s Region (SNNPR), Ethiopia (n=165).

**Table 2:**
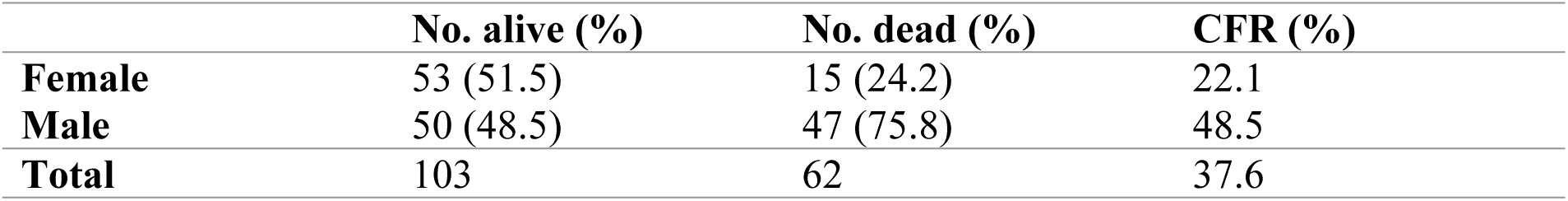
Yellow fever status as of March 2014 by sex, outcome status, and respective case fatality rate (CFR) in Southern Nations, Nationalities, and People’s Region (SNNPR), Ethiopia (n=165).

74.6% of all cases were reported from rural kebeles (villages) in South Ari. Aykamer, Geza and Shepe kebeles contributed 49% of all reported cases and 62% of reported cases within South Ari woreda (Table 1 and Figure 3). Aykamer and Geza kebeles had the highest attack rates (AR) at 103.2 and 104.7 per 10,000 cases respectively. On 10^th^ June, 2013 the SOZ Health Department began an emergency vaccination campaign which targeted 607,462 people (28). HEWs and allied health professionals were able to reach 543,558 people and an overall coverage estimate of 89% was reported from SOZ Health Department in March 2014.

**Figure 3:**
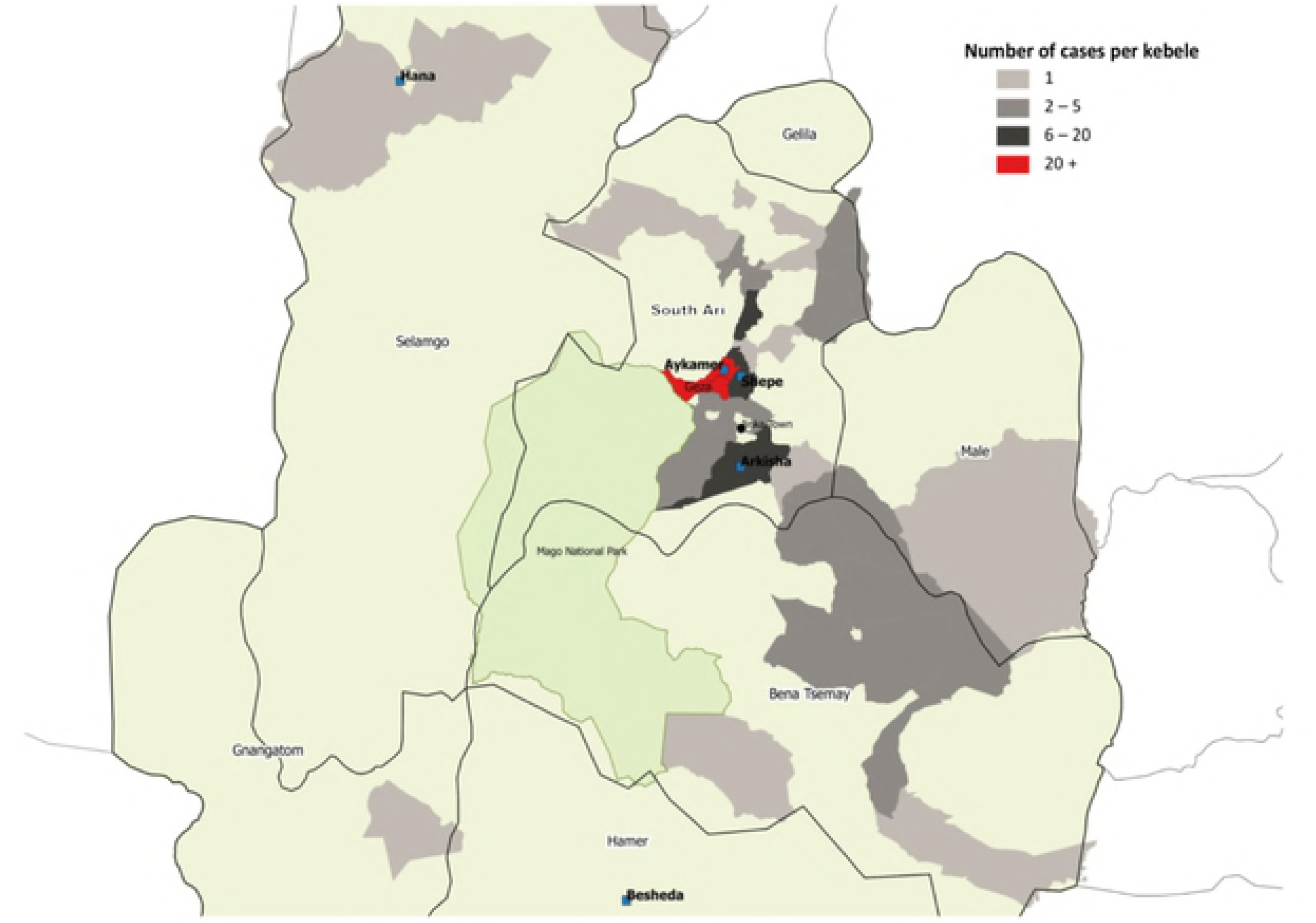
Distribution and frequency of yellow fever cases in South Omo Zone, Ethiopia (n=165).

### Entomological investigation

A total of 688 containers containing water were inspected among 177 households. Overall 240 (34.9%) of the containers were classified as positive, i.e. contained at least one mosquito larva or pupa, across a total of 105 positive households (59.3%). The majority of water-filled containers were found outdoors and filled with rainwater. The locations and types of containers across the kebeles varied, reflecting the differences in larval indices among study sites. No larvae or pupae were found in any water-holding containers in Besheda. A sample of all immature stages were reared to adulthood (Table 3). From Shepe, Arkisha and Aykamer, the main species collected was *Aedes* spp., however from Hana a large proportion was *Culex* spp. A small number of *Toxorhynchites* spp. larvae and pupae were identified in Shepe and Aykamer.

**Table 3:**
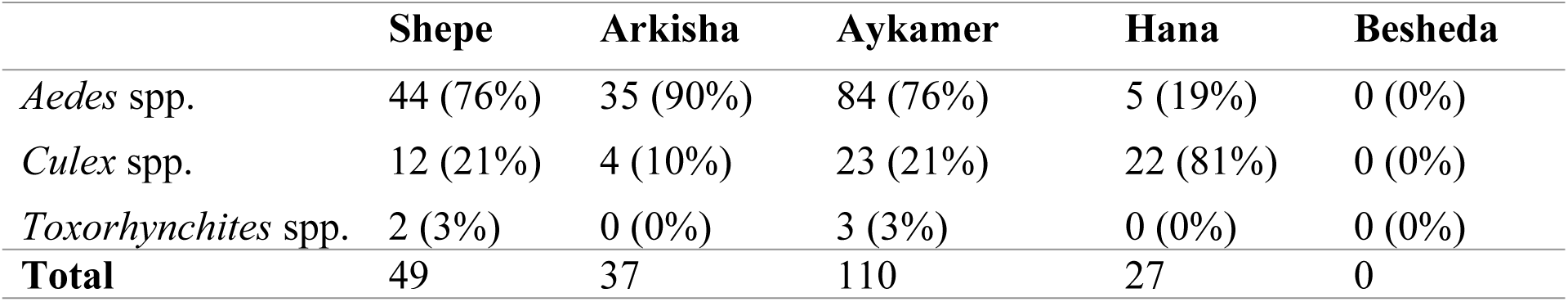
Species of mosquitoes from immature stages reared to adults across the sample sites.

In the highland area of South Ari, the false banana plant is ubiquitously found in close proximity to the home and was the major site for immature *Aedes* spp. stages in Shepe (75% of plants inspected were positive) and Aykamer (64% positive). The second most important breeding sites in these two kebeles were discarded plastic and clay, which were commonly observed outside the home (Figure 4). There was also a high coverage of mixed vegetation close to the home, including sweet potato, maize and false banana, which were often the locations of adult mosquito collections. By comparison, the homes in Besheda were often kept completely clear of any containers, clutter or vegetation. It was very unusual to see any containers outside of the home. In Shepe alone, 102 positive containers were identified, followed by 94 in Aykamer, 24 in Hana, 20 in Arkisha and 0 in Besheda. The types of infested water-holding containers varied between the sample sites depending on the local usage and traditional practices (Table 4).

**Table 4:** Different types of containers inspected, including proportion found positive with either mosquito larvae or pupae, and relative pupal contribution, across the sample kebeles in South Omo Zone, Ethiopia, 2017.

|  | Type of container | No. of containers inspected | No. of positive containers* (%) | No. of pupae (% contribution) |
| --- | --- | --- | --- | --- |
| **Natural breeding sites** | *Ensete ventricosum* (False banana) | 220 | 141 (64.2) | 179 (19.7) |
|  | *Musa* spp. plant (Banana) | 28 | 18 (64.3) | 19 (2.1) |
|  | Tree hole or tree trunk | 6 | 3 (50.0) | 27 (3.0) |
|  | *Colocasia esculenta* (Taro) | 1 | 0 (0.0) | 0 (0.0) |
| **Artificial breeding sites** | Plastic jug | 101 | 5 (4.9) | 3 (0.3) |
|  | Plastic drum / metal drum | 110 | 8 (7.3) | 51 (5.6) |
|  | Discarded plastic | 65 | 14 (21.5) | 86 (9.5) |
|  | Discarded clay pot | 48 | 9 (18.8) | 304 (33.5) |
|  | Clay pot | 45 | 10 (22.2) | 225 (24.8) |
|  | Plastic bowl | 23 | 0 (0.0) | 0 (0.0) |
|  | Metal bowl | 27 | 2 (7.4) | 3 (0.3) |
|  | Discarded tyre | 7 | 4 (57.1) | 8 (0.9) |
|  | Discarded plastic shoe | 4 | 2 (50.0) | 2 (0.2) |
|  | Water pump | 1 | 1 (100.0) | 0 (0.0) |
|  | Discarded metal | 1 | 0 (0.0) | 0 (0.0) |
|  | Glass bottle | 1 | 1 (100.0) | 0 (0.0) |
|  | **Total** | 688 | 34.9 | 907 (100) |

\*presence of at least one larva and/or pupa.

**Figure 4:**
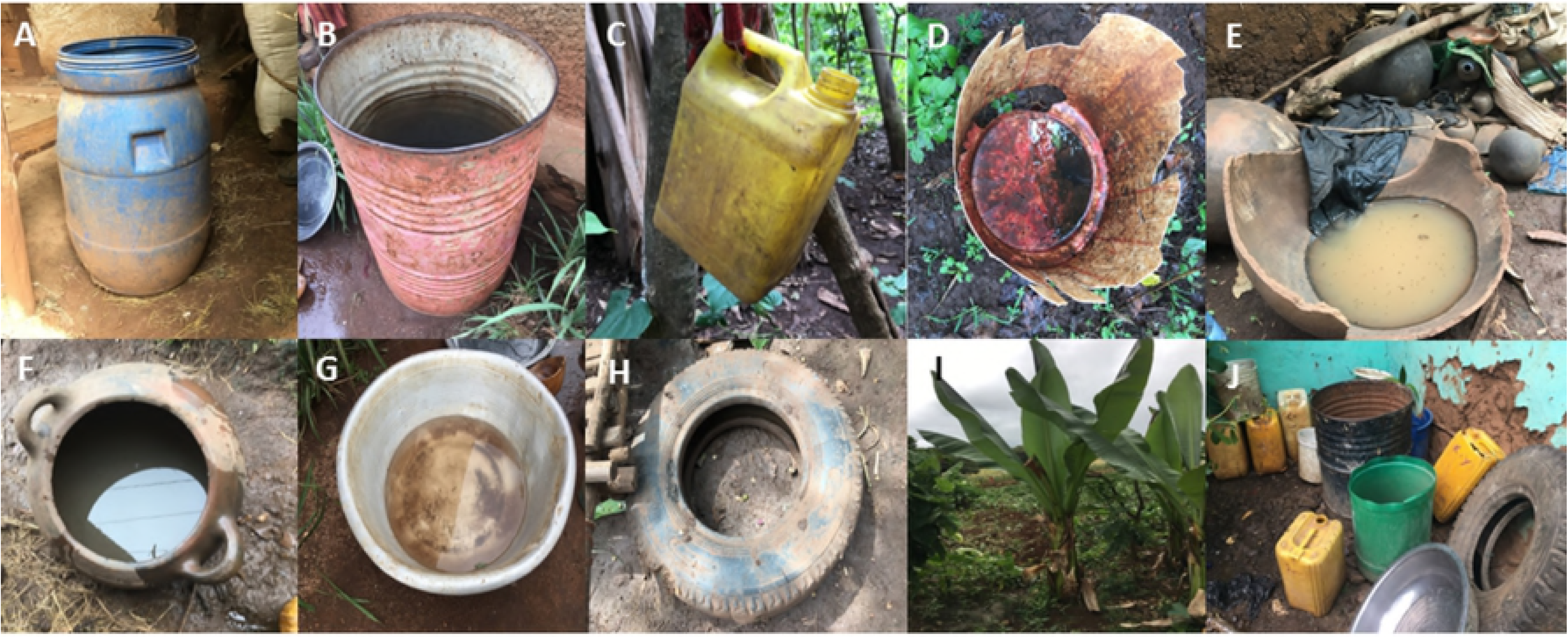
Typical mosquito breeding sites identified in South Omo Zone, Ethiopia, 2017: (a) Plastic drum (b) Metal drum (c) Plastic jug (d) Discarded plastic (e) Discarded clay pot (f) Clay pot (g) Metal bowl (h) Discarded Tyre (i) False banana plant (j) Various discarded items outside the home.

Table 5 shows the Breteau Index (BI), Container Index (CI), Pupal Demographic Index (PDI) and Household Index (HI) for each study site. The highest HIs were recorded in Shepe (79.0%) and Hana (57.1%); both villages were classified as high risk for YFV transmission (WHO threshold of >35%). The highest CIs were in Shepe (57.9%) and Aykamer (75.4%), both of which inferred a high risk (HI>50%). The BIs were higher than the WHO threshold across all sample sites except Besheda (threshold of BI>20); indicating there is no evidence for YFV transmission in Besheda as no immature stage was found and all indices were below the low risk thresholds of HI<4%, CI<3% and BI<5. The PDI was highest in Aykamer (2.24), which also had the highest AR (103.2 cases per 10,000 people) (19). There was no statistically significant correlation between the traditional entomological indices and the AR. However, there was strong positive correlation (*r*=0.9545) between AR and PDI (p=0.0455).

**Table 5:** Distribution of entomological indices and yellow fever (YF) attack rates recorded across the sample kebeles, South Omo Zone, Ethiopia, 2017.

| Woreda | Kebele | Total houses inspected | Household index (%) | Container index (%) | Pupal demographic index* | Breteau Index | YF attack rate^ |
| --- | --- | --- | --- | --- | --- | --- | --- |
| South Ari | Shepe | 43 | 79.0 | 57.9 | 0.48 | 237.2 | 35.5 |
|  | Arkisha | 26 | 38.5 | 18.5 | 0.28 | 76.9 | 9.81 |
|  | Aykamer | 65 | 33.8 | 75.4 | 2.24 | 144.6 | 103.2 |
| Salamago | Hana | 21 | 57.1 | 24.0 | 0.50 | 114.3 | 3.2 |
| Hammer | Besheda | 25 | - | - | - | - | - |
| **Total** |  | **180** | **41.7** | **35.2** | **0.70** | **114.6** | **30.3** |

\*Number of pupae per person.
^Number of cases per 10,000 people.

Screening individual adult female *Aedes* mosquitoes collected using aspiration (n=120 from Shepe, n=12 from Aykamer, n=1 from Arkisha) for YFV and other major arboviruses revealed no evidence of arbovirus infection. Molecular species identification was undertaken by sequencing a fragment of the ITS2 gene for a sub-sample of specimens (approximately 10% of the total number from each location) which included 12 individuals from Shepe, where the majority of adult females were collected, 2 from Aykamer and 1 from Arkisha, in addition to 2 *Ae. bromeliae* specimens from Tanzania for comparison. Figure 5 shows that all specimens collected in our study were phylogenetically similar to GenBank sequences from the *Ae. simpsoni* complex, non-anthropophilic grouping from Mukwaya *et al*. 2000, indicating these specimens are part of the *Ae. simpsoni* complex and group with *Ae. lilii*. Consensus sequences from this study have been submitted to GenBank (Accession numbers MH277621 – MH277635).

**Figure 5:**
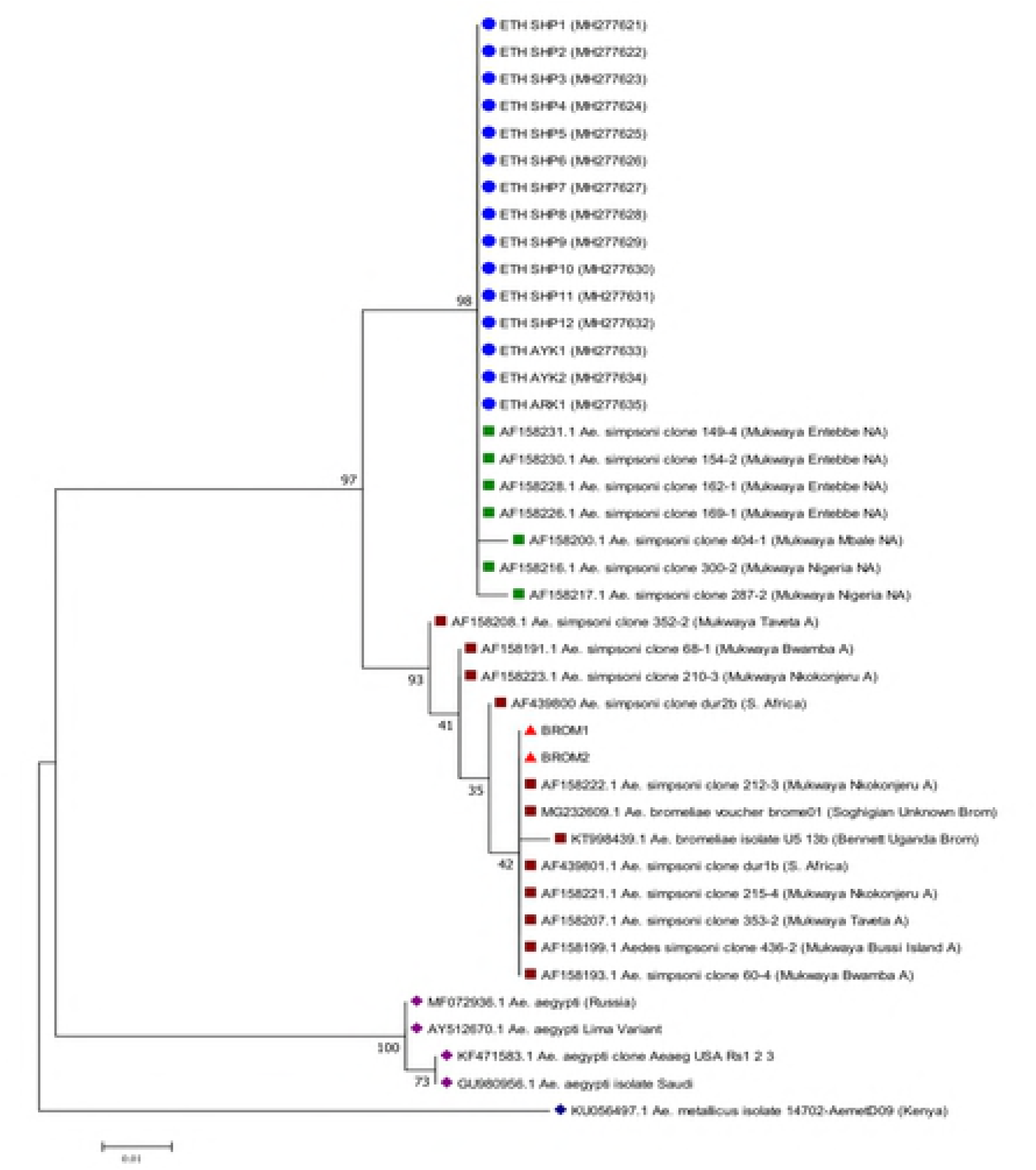
Molecular phylogenetic analysis by Maximum Likelihood method

The evolutionary history was inferred using the Maximum Likelihood method based on the Tamura-Nei model (23). The tree with the highest log likelihood (-587.45) is shown. The percentage of trees in which the associated taxa clustered together is shown next to the branches. Initial tree(s) for the heuristic search were obtained automatically by applying Neighbor-Joining and BioNJ algorithms to a matrix of pairwise distances estimated using the Maximum Composite Likelihood (MCL) approach, and then selecting the topology with superior log likelihood value. The tree is drawn to scale, with branch lengths representing the number of substitutions per site. The analysis involved 41 nucleotide sequences. Codon positions included were 1st+2nd+3rd+Noncoding. All positions containing gaps and missing data were eliminated. There were a total of 230 positions in the final dataset. Evolutionary analyses were conducted in MEGA7(22). Accession numbers for all GenBank sequences included are shown.

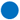 Consensus sequences from field specimens, labelled according to site of collection with newly generated Accession numbers included in brackets (ETH; Ethiopia, SHP; Shepe, AYK; Aykamer, ARK; Arkisha).

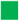 GenBank sequences of *Ae. simpsoni* complex, non-anthropophilic grouping from Mukwaya *et al*. 2000 (29).

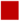 GenBank sequences of *Ae. simpsoni* complex, anthropophilic grouping from Mukwaya *et al*. 2000 (29), in addition to other *Ae. simpsoni* complex and *Ae. bromeliae* GenBank sequences.

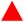 Consensus sequences from *Ae. bromeliae* specimens generated alongside field specimen sequences.

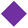 GenBank seqences of *Ae. aegypti*.

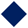 GenBank sequence of *Ae. metallicus*.

### Household and individual characteristics

The KAP respondents comprised of 101 (56.1%) females and 79 (43.9%) males, with 111 (62.3%) between 20-34 years of age (Table 6). The majority either had no education or primary school level (86.6%), while only 5.6% had achieved a higher education level. Among all respondents, 142 (78.9%) were farmers, while the remaining 38 (21.1%) were combinations of other professions.

**Table 6:** Demographic characteristics of study respondents to KAP household questionnaire in South Omo Zone, Ethiopia (n=180).

|  | Variable | Frequency | Percentage % |
| --- | --- | --- | --- |
| Sex | Male | 79 | 43.9 |
|  | Female | 101 | 56.1 |
| Age (years) | <20 | 5 | 2.8 |
|  | 20-24 | 23 | 12.8 |
|  | 25-29 | 39 | 21.7 |
|  | 30-34 | 50 | 27.8 |
|  | 35-39 | 33 | 18.2 |
|  | >40 | 30 | 16.7 |
| Education | None | 96 | 53.3 |
|  | Primary School | 60 | 33.3 |
|  | Secondary School | 14 | 7.8 |
|  | Higher Education | 10 | 5.6 |
| Marital Status | Married | 152 | 84.4 |
|  | Single | 12 | 6.7 |
|  | Divorced | 2 | 1.1 |
|  | Widowed | 14 | 7.8 |
| **Occupation** | Unemployed | 14 | 7.8 |
|  | Farmer | 142 | 78.9 |
|  | Shopkeeper | 1 | 0.6 |
|  | Forest-worker | 1 | 0.6 |
|  | Trader | 4 | 2.2 |
|  | Teacher | 3 | 1.6 |
|  | Student | 6 | 3.3 |
|  | Other | 9 | 5.0 |
| **No. of people in the household** | <4 | 26 | 14.4 |
|  | 4 | 43 | 23.9 |
|  | 5 | 29 | 16.1 |
|  | 6 | 40 | 22.2 |
|  | 7 | 19 | 10.6 |
|  | >7 | 23 | 12.8 |

### Knowledge of YF

158 (87.8%) of the respondents had previously heard of YF, with the majority able to identify the general symptoms of YF, including fever (82.2%) and headache (82.2%) (Table 7). However, fewer participants were able to identify more specific YF signs and symptoms, including jaundice (61.7%), muscle pain (74.5%) and bloody vomiting (56.1%). Most participants stated that mosquitoes transmitted YFV (82.8%) (Table 7). However, knowledge levels were lower concerning which mosquito species transmitted the diseases, with only 54 (30%) aware that it was different species to those capable of transmitting malaria. Many respondents (57.2%) affirmed that YFV could be transmitted through ordinary person to person contact, although knowledge was higher that it is not possible to transmit YFV via food or water (62.8%). Many thought YF mosquitoes were most likely to bite at night time (58.9%). The majority of respondents knew that mosquitoes could breed in standing water (83.9%) but much fewer were aware that this was possible inside the home (46.7%). 138 (76.7%) agreed that removing or covering standing water helps to prevent mosquito breeding and 145 (80.6%) agreed that pouring chemical into standing water was effective at killing mosquito larvae.

**Table 7:** Knowledge of yellow fever (YF) signs, symptoms and transmission modes among study respondents in South Omo Zone, Ethiopia, 2017 (n=180).

| **KNOWLEDGE** | **Frequency** | **Percentage %** |
| --- | --- | --- |
| **Have you heard of yellow fever?** |  |  |
| Yes | 158 | 87.8 |
| No | 22 | 12.2 |
| **SIGNS AND SYMPTOMS** |  |  |
| **Is fever a symptom of YF?** |  |  |
| Yes | 148 | 82.2 |
| No | 8 | 4.5 |
| Don't know | 2 | 1.1 |
| No answer | 22 | 12.2 |
| **Is headache a symptom of YF?** |  |  |
| Yes | 148 | 82.2 |
| No | 8 | 4.5 |
| **Don't know** | 2 | 1.1 |
| **No answer** | 22 | 12.2 |
| **Is jaundice a symptom of yellow fever?** |  |  |
| **Yes** | 111 | 61.7 |
| **No** | 37 | 20.5 |
| **Don't know** | 10 | 5.6 |
| **No answer** | 22 | 12.2 |
| **Is muscle pain a symptom of YF?** |  |  |
| **Yes** | 134 | 74.5 |
| **No** | 20 | 11.1 |
| **Don't know** | 4 | 2.2 |
| **No answer** | 22 | 12.2 |
| **Is rash a symptom of YF?** |  |  |
| **Yes** | 97 | 53.9 |
| **No** | 50 | 27.8 |
| **Don't know** | 11 | 6.1 |
| **No answer** | 22 | 12.2 |
| **Is bloody vomiting a symptom of YF?** |  |  |
| **Yes** | 101 | 56.1 |
| **No** | 29 | 16.1 |
| **Don't know** | 28 | 15.6 |
| **No answer** | 22 | 12.2 |

| **TRANSMISSION** | **Frequency** | **Percentage %** |
| --- | --- | --- |
| **Do mosquitoes transmit YF?** |  |  |
| **Yes** | 149 | 82.8 |
| **No** | 9 | 5.0 |
| **Don't know** | 3 | 1.7 |
| **No answer** | 19 | 10.5 |
| **Are they the same mosquitoes that transmit malaria?** |  |  |
| **Yes** | 81 | 45.0 |
| **No** | 54 | 30.0 |
| **Don't know** | 26 | 14.4 |
| **No answer** | 19 | 10.6 |
| **Does ordinary person to person contact transmit YF?** |  |  |
| **Yes** | 103 | 57.2 |
| **No** | 46 | 25.6 |
| **Don't know** | 12 | 6.6 |
| **No answer** | 19 | 10.6 |
| **Is YF transmitted through food and water?** |  |  |
| **Yes** | 38 | 21.1 |
| **No** | 113 | 62.8 |
| **Don't know** | 10 | 5.5 |
| **No answer** | 19 | 10.6 |
| **When are the YF mosquitoes most likely to bite?** |  |  |
| **Night** | 106 | 58.9 |
| **Day** | 16 | 8.9 |
| **Both** | 39 | 21.6 |
| **Don't know** | 19 | 10.6 |
| **Do the YF mosquitoes breed in standing water?** |  |  |
| **Yes** | 151 | 83.9 |
| **No** | 2 | 1.1 |
| **Don't know** | 8 | 4.4 |
| **No answer** | 19 | 10.6 |
| **Can mosquitoes breed inside the home?** |  |  |
| **Yes** | 84 | 46.7 |
| **No** | 64 | 35.6 |
| **Don't know** | 13 | 7.1 |
| **No answer** | 19 | 10.6 |
| **Does removal or covering of standing water prevent mosquito breeding?** |  |  |
| **Yes** | 138 | 76.7 |
| **No** | 10 | 5.5 |
| **Don't know** | 13 | 7.2 |
| **No answer** | 19 | 10.6 |
| **Can pouring chemical into standing water kill mosquito larvae?** |  |  |
| **Yes** | 145 | 80.6 |
| **No** | 6 | 3.3 |
| **Don't know** | 10 | 5.5 |
| **No answer** | 19 | 10.6 |

### Attitudes towards prevention and control of YF

147 (81.7%) of study respondents stated that YF is a serious illness, with the reasons for being so including bloody vomiting, high CFR, and becoming more dangerous without early treatment (Table 8). Respondents agreed that both controlling breeding sites of mosquitoes (87.2%) and vaccination (75.6%) were good strategies to prevent YF, and that communities have a role to play in controlling these mosquitoes (86.7%). Fewer participants (56.1%) thought that it was the health post’s responsibility to prevent YF.

**Table 8:**
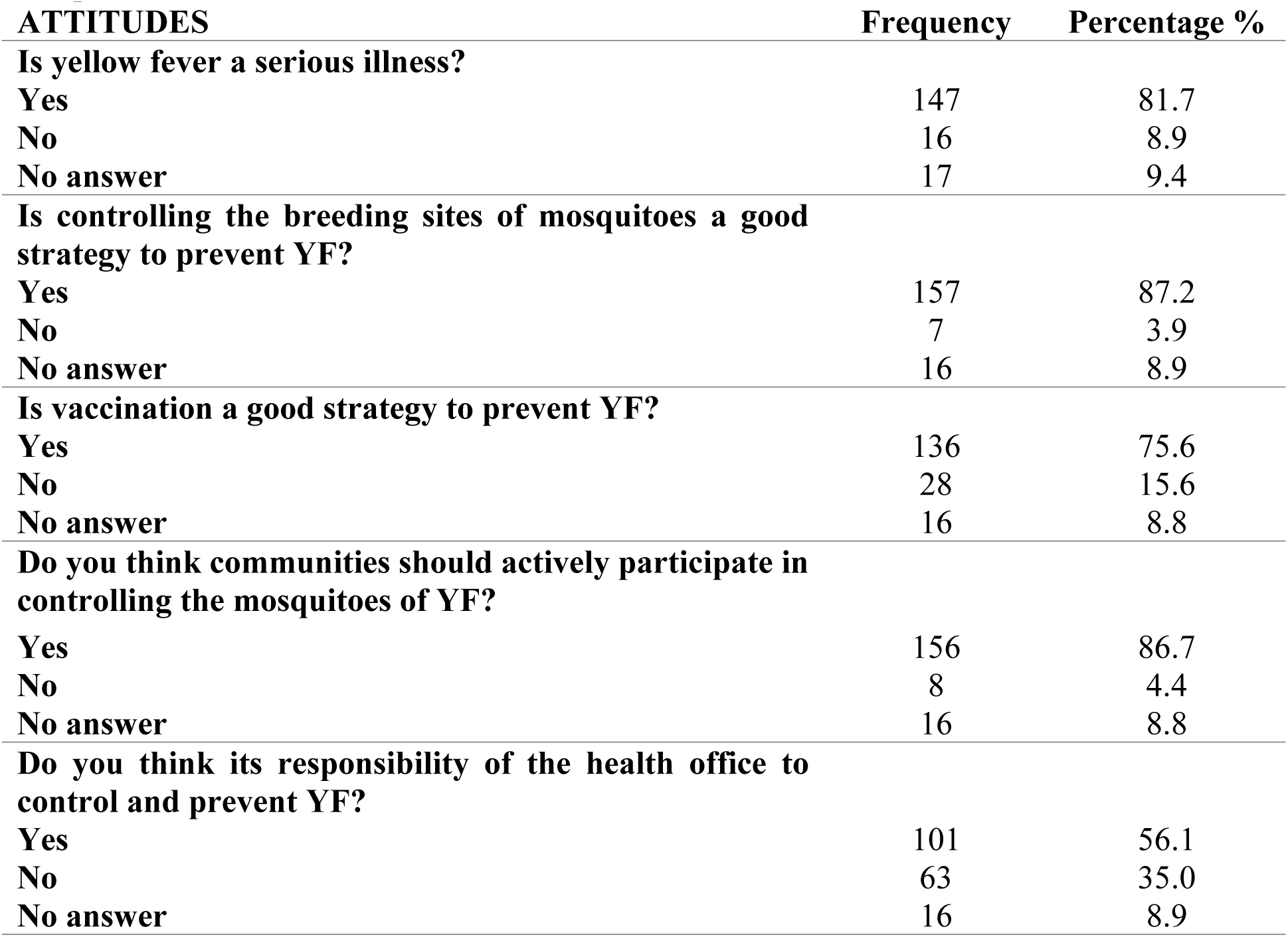
Attitudes of study respondents towards yellow fever (YF) in South Omo Zone, Ethiopia, 2017 (n=180).

| **ATTITUDES** | **Frequency** | **Percentage %** |
| --- | --- | --- |
| **Is yellow fever a serious illness?** |  |  |
| **Yes** | 147 | 81.7 |
| **No** | 16 | 8.9 |
| **No answer** | 17 | 9.4 |
| **Is controlling the breeding sites of mosquitoes a good strategy to prevent YF?** |  |  |
| **Yes** | 157 | 87.2 |
| **No** | 7 | 3.9 |
| **No answer** | 16 | 8.9 |
| **Is vaccination a good strategy to prevent YF?** |  |  |
| **Yes** | 136 | 75.6 |
| **No** | 28 | 15.6 |
| **No answer** | 16 | 8.8 |
| **Do you think communities should actively participate in controlling the mosquitoes of YF?** |  |  |
| Yes | 156 | 86.7 |
| No | 8 | 4.4 |
| No answer | 16 | 8.8 |
| **Do you think its responsibility of the health office to control and prevent YF?** |  |  |
| Yes | 101 | 56.1 |
| No | 63 | 35.0 |
| No answer | 16 | 8.9 |

### Practices regarding YF prevention

166 (92%) respondents self-reported actively reducing mosquitoes near their homes, through preventing standing water (89.4%), using insecticide-treated nets in the home (85.6%), using smoke to drive mosquitoes away (86.7%) and wearing clothes to prevent mosquito bites (92.8%) (Table 9). 145 (81%) respondents reported that the government had come to their houses to spray insecticides on their walls in the past. The majority of respondents reported covering water containers in the home (90.0%), with many claiming to clean their water filled containers every day (37.3%) or once a week (51.7%). Most respondents reported turning their containers upside down to avoid water collection (84.4%).

**Table 9:** Practices for yellow fever (YF) prevention and mosquito control of study respondents in South Omo Zone, Ethiopia, 2017 (n=180).

| **PREVENTING MOSQUITO-MAN CONTACT** | **Frequency** | **Percentage %** |
| --- | --- | --- |
| **Do you do anything to reduce mosquitoes?** |  |  |
| Yes | 166 | 92.2 |
| No | 14 | 7.8 |
| **Has the government come to spray insecticide to reduce mosquitoes?** |  |  |
| Yes | 145 | 80.6 |
| No | 35 | 19.4 |
| **Do you prevent standing water around the house to reduce mosquitoes?** |  |  |
| Yes | 161 | 89.4 |
| No | 19 | 10.6 |
| **Do you use insecticide treated nets to protect against mosquitoes in the home?** |  |  |
| Yes | 154 | 85.6 |
| No | 26 | 14.4 |
| **Do you use smoke to drive mosquitoes away?** |  |  |
| Yes | 156 | 86.7 |
| No | 24 | 13.3 |
| **Do you cover your body with clothes to prevent against mosquito bites?** |  |  |
| Yes | 167 | 92.8 |
| No | 13 | 7.2 |
| **ELIMINATING BREEDING SITES** |  |  |
| **Do you cover water containers in the home?** |  |  |
| Yes | 162 | 90.0 |
| No | 18 | 10.0 |
| **How often do you clean water filled containers and ditches around the house?** |  |  |
| Everyday | 67 | 37.3 |
| Once a week | 93 | 51.7 |
| Once a month | 9 | 5.0 |
| Never | 7 | 3.9 |
| No answer | 4 | 2.1 |
| **Do you turn containers upside down to avoid water collection?** |  |  |
| Yes | 152 | 84.4 |
| No | 27 | 15.0 |
| No answer | 1 | 0.6 |

### Sources of information about YF

The majority of respondents (76.8%) reported receiving information solely from their HEWs, particularly as most live in rural kebeles without access to radio or TV. A large proportion of respondents knew of neighbours or family members who had YF in the past, with many reporting cases that occurred 5-6 years ago. However, individuals often cited cases occurring between 2014 and 2017, particularly in Arkisha where 64% of respondents reported knowing someone who had YF in 2016 or 2017. 60% of respondents from Arkisha self-reported having suffered from YF themselves, while no one in Besheda reported any cases of YF for either themselves or anyone else they knew.

Self-reported vaccination history, collected alongside household KAP survey data, reported only 62% of respondents having been vaccinated in the YF campaign in 2013, compared to the 89% coverage estimated by SOZ Health Department by March 2014. All of those who reported being vaccinated, had done so at their kebele health post in 2013.

21 (11.7%) study respondents reported working in Mago National Park or other forest areas, particularly from Arkisha which borders the park. 45 (25%) people reported having contact with monkeys near to their homes. From the rural kebeles, little migration had occurred, however, in Hana, almost every respondent (90.5%) had moved to the region for work, in particular to work at the Omo Kuraz Sugar Factory project, or to serve this growing urban community.

### Predictors for knowledge, attitudes and practices

Regarding KAP scores, 53.3% of participants achieved a good (at least 75%) knowledge score, 78.9% achieved a good attitude score and 70.6% a good practices score. In the univariate analysis of the association between KAP score and a number of independent variables (including socio-economic variables, past YFV infection and YF vaccination status), there was increased odds of having good knowledge if the respondent lived in Arkisha (OR:13.88; 95% CI 4.18-46.02), Aykamer (OR: 18.89; 95% CI 6.74-52.94), Hana (OR: 7.40; 95% CI 2.22-24.65) and Besheda (OR:5.22; 95% CI1.61-16.95), compared to a respondent living in Shepe (Table 10).

**Table 10:**
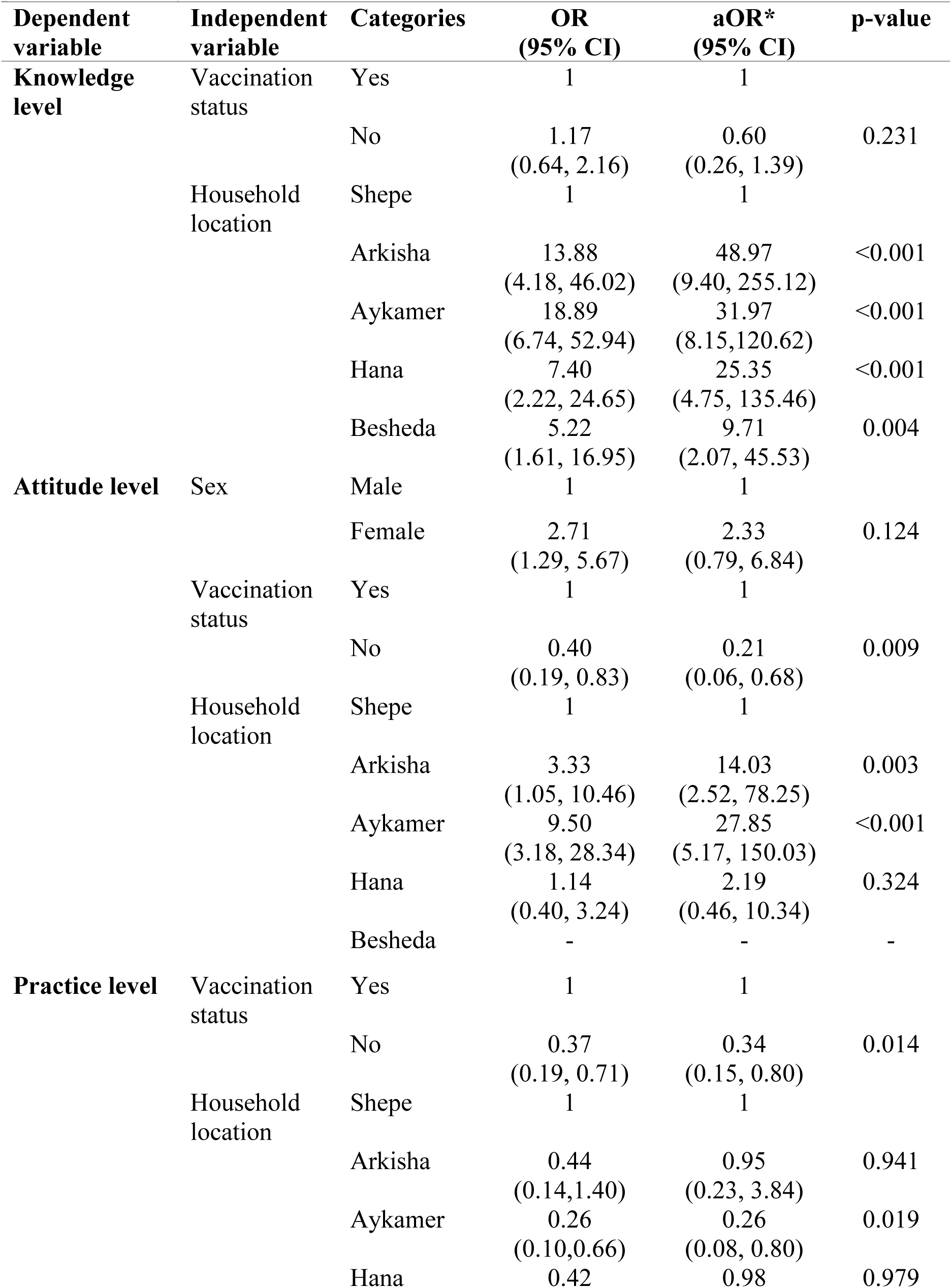
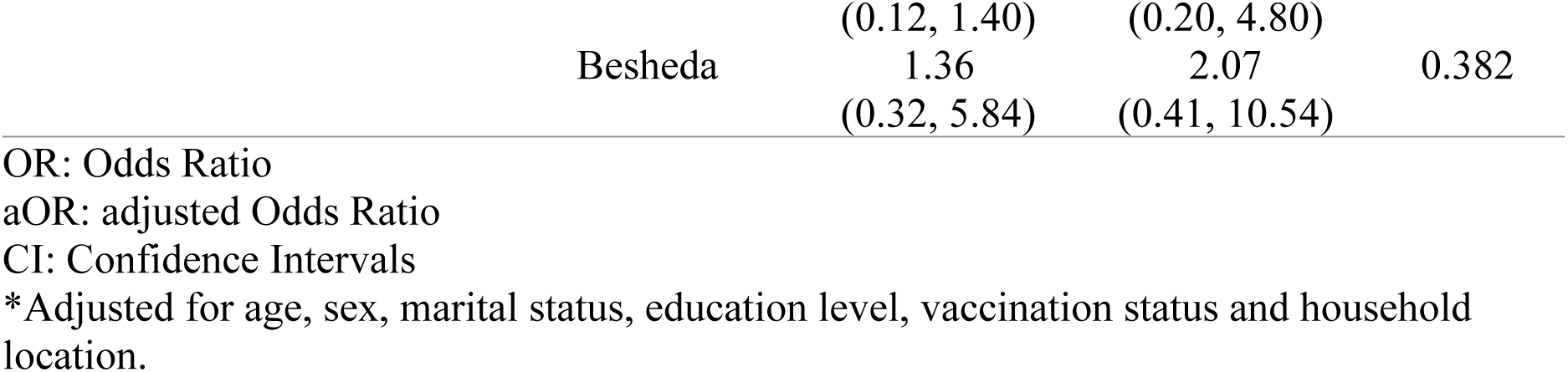
Multivariate logistic regression showing predictors of knowledge, attitude and practices levels (good *vs*. bad).

| Dependent variable | Independent variable | Categories | OR (95% CI) | aOR* (95% CI) | p-value |
| --- | --- | --- | --- | --- | --- |
| Knowledge level | Vaccination status | Yes | 1 | 1 |  |
|  |  | No | 1.17<br>(0.64, 2.16) | 0.60<br>(0.26, 1.39) | 0.231 |
|  | Household location | Shepe | 1 | 1 |  |
|  |  | Arkisha | 13.88<br>(4.18, 46.02) | 48.97<br>(9.40, 255.12) | <0.001 |
|  |  | Aykamer | 18.89<br>(6.74, 52.94) | 31.97<br>(8.15, 120.62) | <0.001 |
|  |  | Hana | 7.40<br>(2.22, 24.65) | 25.35<br>(4.75, 135.46) | <0.001 |
|  |  | Besheda | 5.22<br>(1.61, 16.95) | 9.71<br>(2.07, 45.53) | 0.004 |
| Attitude level | Sex | Male | 1 | 1 |  |
|  |  | Female | 2.71<br>(1.29, 5.67) | 2.33<br>(0.79, 6.84) | 0.124 |
|  | Vaccination status | Yes | 1 | 1 |  |
|  |  | No | 0.40<br>(0.19, 0.83) | 0.21<br>(0.06, 0.68) | 0.009 |
|  | Household location | Shepe | 1 | 1 |  |
|  |  | Arkisha | 3.33<br>(1.05, 10.46) | 14.03<br>(2.52, 78.25) | 0.003 |
|  |  | Aykamer | 9.50<br>(3.18, 28.34) | 27.85<br>(5.17, 150.03) | <0.001 |
|  |  | Hana | 1.14<br>(0.40, 3.24) | 2.19<br>(0.46, 10.34) | 0.324 |
|  |  | Besheda | - | - | - |
| Practice level | Vaccination status | Yes | 1 | 1 |  |
|  |  | No | 0.37<br>(0.19, 0.71) | 0.34<br>(0.15, 0.80) | 0.014 |
|  | Household location | Shepe | 1 | 1 |  |
|  |  | Arkisha | 0.44<br>(0.14, 1.40) | 0.95<br>(0.23, 3.84) | 0.941 |
|  |  | Aykamer | 0.26<br>(0.10, 0.66) | 0.26<br>(0.08, 0.80) | 0.019 |
|  |  | Hana | 0.42 | 0.98 | 0.979 |

|  |  |  |  |
| --- | --- | --- | --- |
|  | (0.12, 1.40) | (0.20, 4.80) |  |
| Besheda | 1.36 | 2.07 | 0.382 |
|  | (0.32, 5.84) | (0.41, 10.54) |  |

OR: Odds Ratio
aOR: adjusted Odds Ratio
CI: Confidence Intervals
\*Adjusted for age, sex, marital status, education level, vaccination status and household location.

This analysis also predicted increased odds of having a good attitude score if the respondent was female (OR: 2.71; 95% 1.29-5.67), and by household location, with a decreased odds of good attitude by lack of YF vaccination (OR: 0.40, 95% CI 0.19-0.83) (Table 10). A good practice level score was also found to be predicted by YF vaccination status (OR: 0.37, 95% 0.19-0.71). After adjusting for potential confounders in a multivariate analysis, sex was no longer found to be a significant predictor of attitude levels; or vaccination status on knowledge level (Table 10).

The correlation of KAP scores revealed a slight positive correlation between knowledge and attitude scores (*r*=0.41, p<0.001); while no significant correlation was found between knowledge and practices (p=0.373), or attitude and practices (p=0.471) (Table 11).

**Table 11:** Correlation between knowledge, attitude and practices scores.

| Variables | Correlation co-efficient (*r*) | p-value |
| --- | --- | --- |
| Knowledge-Attitudes | 0.41 (0.24, 0.58) | <0.001 |
| Knowledge-Practices | -0.07 (-0.23, 0.09) | 0.373 |
| Attitudes-Practices | 0.05 (-0.08, 0.18) | 0.471 |

## Discussion

To reduce the risk of future YFV and other arboviral disease outbreaks, it is important to understand past outbreaks and community perceptions and preventative practices in at-risk areas, to guide appropriate interventions and policy recommendations. From 2012-2014, 165 cases were reported in SOZ, including 62 fatalities. YF was unfamiliar to health professionals at this time as it was an unexpected outbreak of a disease that had not been seen locally since 1962 (30). With the majority of cases identified in rural kebeles, it is highly likely that the prevalence and severity of the outbreak was under-reported.

The epidemiological data indicate that the case number was higher in males, which is in line with previous YF outbreaks, such as the 1960 outbreak in Ethiopia which had a male:female case ratio of 1.6:1 (31) as well as in Uganda (2011) where the sex-specific AR was 16.5 (male) *vs*. 9.6 (female) per 100,000 (32). In SOZ, the CFR in males (48.5%) was almost double compared to females (22.1%), which was consistent with the YF outbreak in Uganda (29.6% in males compared to 17.8% in females) and may be explained by males often reporting later (and sicker) to health facilities, due to underestimating the severity of the disease at symptom onset. By contrast, during outbreaks in The Gambia (1978-1979) and Ghana (1969-1970), children <15 years of age were the most affected, due to the stopping of routine mass YF vaccination campaigns in the early 1960s (33).

The majority of cases were seen in 15-44 year olds (75.8%) who may have been infected while working outdoors during the day and dusk, the peak mosquito biting times; a risk factor also described in Kenya (1992-1993) where 81% of cases were <40 years old (34). The overall CFR in SOZ was 37.6%, which is higher than in the Darfur epidemic of 2012 (20.3%) and in Uganda in 2011 (24.9%) (33). Only 4 cases were reported from outside of SOZ, which may be from individuals visiting the region, e.g. becoming infected at the weekly Saturday market in Jinka Town; rather than autochthonous YFV transmission. However, with reports of *Aedes* spp. in Gamo Gofa (14), the potential for local transmission is possible (as seen in 1966, a region previously unaffected during the 1960-1962 outbreak). Areas outside of SOZ were not included in the mass vaccination campaign of 2013, leaving this population immunologically-vulnerable to future outbreaks.

The mass emergency vaccination campaign of 2013 appeared to curb the outbreak as the number of cases declined post-campaign and a high administrative vaccination coverage was reported. However, cases were still appearing in early 2014: for example, 3 cases in Malle woreda (previously unaffected during the peak period of the outbreak) were detected in individuals who had not been vaccinated (35), and there are informal reports from South Omo and Arba Minch of recent YF cases; highlighting the urgent need for a follow up vaccination campaign and strengthened case investigation. The peak of the reported cases occurred in April and May of 2013, which could be explained by the hot and wet climate in the region during this time, encouraging mosquito breeding. However, this also coincided with the YFV laboratory confirmation which increased disease awareness among community and health professionals.

In early 2014, the EPHI conducted a training for health managers at regional and district levels on YF. However, by this time, the peak of the outbreak was already over. In addition, surveillance was slow and took almost 8 months to confirm the outbreak. These delays were due to a combination of a lack of in-country laboratory diagnostics (requiring samples to be sent to the Regional Reference Laboratory at Institut Pasteur, Dakar for analysis), non-specificity of the case definition, and misdiagnosis by clinicians (16)(36). Through informal discussions, it was noted that most health professionals (particularly doctors) working in Jinka Hospital rotate to new hospitals after 2-3 years. Many of the doctors currently at the hospital did not know the signs, symptoms or treatment for YF, as no follow up trainings by the EPHI have taken place for newer recruits. Therefore, knowledge levels of healthcare professionals can be assumed low in the case of a future outbreak.

During the entomological investigations, a total of 688 artificial and natural containers were inspected from across 177 households. Most of the containers were found outside of the home, and due to the ongoing rains in South Ari, filled with rainwater. None of the kebeles sampled had piped water, so buckets and drums were often used for water storage. The urban area of Hana served a larger number of people per household, and therefore stored a larger proportion of water, increasing the number of potential sites for harbouring immature stages.

The main breeding site in Aykamer and Shepe was the false banana plant, a crop found ubiquitously in South Ari. Traditional practices ensure the plant is cultivated close to the home, particularly as it requires a larger amount of dung and nutrients than the regular banana plant. The plant has multiple purposes, including its leaves which are used to transport cabbage to market and its stem and roots are used as food (the “false banana” that fruits is inedible). In addition, the plant grows for about 4 years, in comparison to the banana plant which grows, fruits and dies within a few weeks, providing a longer life-span to act as a mosquito breeding site. In the hotter and drier months, the water in the large plant stem has previously been reported to resist evaporation, making mosquito breeding possible all year round (30). A recent mini-drought in Arkisha resulted in the loss of all false banana plants, however, usually it is pervasive in this kebele. Therefore, breeding sites described in Arkisha during this study may not be entirely representative. In addition, previous studies identified the taro plant as an important breeding site for *Aedes* spp. in East Africa, however, in this study it was often difficult to isolate water from the plant and only a small number were inspected, therefore its significance is most probably under-estimated (30).

The greatest pupal contribution across the study sites was from clay pots (both used and discarded) as well as the false banana plant. The clay pots were used for multiple purposes, but commonly found to store water during house construction or those that were broken and discarded containing rainwater; in the former, as the water was not being consumed, it was left uncovered. Drums were also often used for water storage; however, most were found negative for mosquito larvae. One study in Dire Dawa, Ethiopia, found that artificial breeding sites (in particular tyres and plastic drums) were the primary source of vectors (37,38), however there sampling was conducted in an urban area. Various studies found differing types of containers responsible for *Aedes* spp. breeding, such as discarded tyres in Tanzania (39), medium storage containers in Nigeria (40) and natural sites in Kenya (34,41).

Larval entomological surveys were conducted to understand the larval density and mosquito abundance, to determine the future risk of YFV transmission. In terms of entomological risk indices, all kebeles except Besheda were above the WHO high risk thresholds for one or more entomological index, indicating evidence for future local YFV transmission. As previous studies have reported a number of limitations associated with measuring larval indices, pupal numbers per person were also calculated (PDI) (42). The PDI was found highest in Aykamer (2.24), with a strong positive correlation between the AR and PDI (*r*=0.9545, p=0.0455), and therefore it can be considered a significant predictor of YF risk. A higher PDI was also observed in rural areas, consistent with a previous study of *Ae. aegypti* breeding sites in Kenya (43).

The ITS2 sequencing data from adult *Aedes* captured in and around households in this study, in comparison with currently available sequences, indicated that they were most closely related to non-anthropophilic members of the *Ae. simpsoni* complex from Uganda and Nigeria (29). This complex comprises three known species including *Ae. simpsoni* s.s. *Ae. lilii* and *Ae. bromeliae*, of which the latter is considered anthropophilic and has previously been incriminated as the principle vector of YF epidemics in Ethiopia (16,44)). Adult mosquitoes collected from SOZ were most likely *Ae. lilii* (*Ae. simpsoni* s.s. is confined to southern Africa) and therefore the lack of detection of any medically important arboviruses which are transmitted through human blood-feeding would be expected in this species given no females have been recorded biting humans (29). While *Ae. bromeliae* can breed across varied ecologies, *Ae. lilii* has been sampled from a more restricted range of plant axils (*Musa* spp. *Colocasia* spp. *Dracena* spp. and *Sansevieria* spp.) and not previously from inside the domestic environment (45,46). Additional sampling efforts are warranted to identify the vector species responsible for YFV transmission and its breeding patterns, to further define the local distribution and ecology of *Ae. lilii*, and in particular, the host feeding behaviour of the species in this specific locality.

The KAP survey was completed by an equal split of male and female respondents (43.9% and 56.1% respectively) and the majority did not have or had very little education. Despite this, good knowledge (53.3%), attitude (78.89%) and practice (70.56%) scores were surprisingly high across all respondents, which may also be explained by the confusion with, and fear of, malaria. The results from the study showed that 87.8% of respondents had heard of yellow fever (Amharic translation: *bicha woba*). However, the direct translation of *bicha* = yellow and *woba* = malaria, meant the direct translation of *bicha woba* was “yellow malaria” which resulted in a large amount of confusion between YF and severe malaria. Therefore, if respondents thought the questionnaire was about severe malaria, information bias would result in an overestimation of YF knowledge levels. Knowledge levels of general symptoms were high, while the more specific symptoms were much lower. 57.2% of people thought YF could be transmitted from person to person, which could be an issue when people are seeking treatment and likewise when administering care to others. This finding is in agreement with a recent study by Legesse *et al*. (2018) that also found a fairly high percentage (55.9%) of respondents in SOZ believed YFV could be transmitted through breathing (47).

There were high knowledge levels that YFV was transmitted by a mosquito (82.8%), in contrast to Legesse *et al*. (2018) who found only 37.6% knew YFV was vector-borne. However, because awareness of malaria transmission by mosquitoes was high, the true knowledge level may have been overestimated in our study. Knowledge on malaria was not quantitatively assessed in this study, however, studies by Abate *et al*. (2013) in other parts of Ethiopia showed 85% of respondents to a malaria KAP survey stated its transmission to be by mosquitoes (48). Only 30.5% knew that the mosquito could bite during the daytime and therefore, many people may not be adequately protecting themselves from mosquito bites. 81.7% stated that YF was a serious illness, with reasons including bloody vomiting and problems with the brain, which is also in agreement with the confusion over YF and severe and/or cerebral malaria.

A number of respondents from Arkisha reported knowing someone who had suffered from YF in 2016 or 2017, or self-reporting having suffered from YF themselves. This is consistent with the reporting of 22 suspected cases in Arkisha (sub-cluster: Giste) in March 2016 by PHEM (49). However, according to a Weekly Humanitarian Bulletin for Ethiopia in June 2016, these cases turned out to be negative (50).

The only significant determinants of KAP scores were household location for knowledge score; vaccination status and household location for attitude score; and vaccination status for practice scores. As the majority of those vaccinated would have been so in their local kebele health post by a HEW, they would have also been educated on the disease at this time. Loudspeaker announcements throughout the kebele told heads of households to come and get their families vaccinated, rather than actively seeking individuals for vaccination. This suggests those that were the most active in ensuring the health of their household (measured by self-reported vaccination status), had the highest KAP levels of YF.

In addition, the finding of household location being predictive of the KAP scores was in agreement with how the respondents gathered their information on YF, with 76.8% reporting HEWs as their only source. This suggests the odds of having good knowledge; which were 49.0 higher in Arkisha, 32.0 in Aykamer, 25.4 in Hana and 9.7 in Besheda compared to Shepe; could be explained by how active and knowledgeable the HEWs were in the respective localities. However, this may also have been influenced by interviewer bias. The same was seen for attitude levels, with the odds of having a good score higher in other kebeles, compared to Shepe.

### Study limitations

There are several weaknesses in the reported study design, which need to be considered when interpreting the data. The line list of YF cases from the PHEM contained missing data on symptoms, occupation, travel history or past vaccination history, and therefore, a clinical epidemiological analysis was not conducted and the scope for YF risk factor analysis was limited. Regarding entomological indices, cross-sectional sampling was undertaken and thus it was not possible to assess the relative importance of each breeding site over time or with respect to the rainy season; nor was it logistically feasible to extend mosquito collections to the forest, to improve our understanding of local sylvatic YFV transmission. It was also not possible to retrieve historical climatic data during the YFV outbreak period, which could have helped to identify environmental risk factors for this outbreak. Due to the relatively low numbers of mosquito species collected, it was not possible to identify the vector species responsible for YFV transmission nor successfully detect any arboviruses within sampled mosquitoes. Migrant workers and nomadic populations may not have been captured within this study due to the time of day in which the questionnaires were completed, and therefore a follow up survey should be conducted, also including those working for the Omo Kuraz Sugar Factory in Salamago. All interviews were conducted by the HEWs in the language most comfortable to the interviewee. However, in each kebele we worked with different HEWs to ensure community acceptability, but this may have introduced interview bias when completing the questionnaire, translating and interpreting answers. Due to small numbers of participants from each kebele, data were pooled for analysis; however, this may have concealed variations in KAP level by location, therefore multivariate analysis was performed to account for this. Finally, self-reporting of vaccination status also introduced an unascertainable amount of recall bias, but unfortunately, vaccination cards were not provided during the mass campaign.

### Conclusions

Despite YFV re-emerging in recent years, little research has been conducted across endemic countries to understand the risk for further outbreaks. This study suggests that following an outbreak in 2012-2014, the SOZ area in southwestern Ethiopia still remains at risk for future YFV transmission. With the high density of *Aedes* mosquitoes breeding close to human habitats, and relatively low community knowledge levels of YF prevention methods, study findings highlight the importance of providing information to these at-risk communities. Further research needs to be conducted in the surrounding areas, to identify the major vector species of YFV, to understand the local sylvatic YFV transmission in forest areas and to further define the local distribution of *Ae. simpsoni* complex mosquitoes. Follow-up vaccination campaigns should be considered to target remaining pockets of potential unvaccinated populations, in parallel to introduction of the vaccination into the country’s national childhood immunization regimen. Future training of HEWs and health professionals is necessary to ensure sustained high knowledge of both professionals and the community. Overall, study results emphasise the need to strengthen local and national disease surveillance and in-country laboratory capacity to ensure more rapid detection and response to future outbreaks.

## Abbreviations

Attack Rate (AR)
Breteau Index (BI)
Case Fatality Rate (CFR)
Chikungunya virus (CHIKV)
Container Index (CI)
Dengue virus (DENV)
Eliminating Yellow Fever Epidemics (EYE)
Ethiopian Public Health Institute (EPHI)
Health Extension Worker (HEW)
Household Index (HI)
Knowledge, Attitudes and Practices (KAP)
Public Health Emergency Management (PHEM)
Pupal Demographic Index (PDI)
Rift Valley fever virus (RVFV)
South Omo Zone (SOZ)
Southern Nations, Nationalities, and People’s Region (SNNPR)
West Nile virus (WNV)
World Health Organization (WHO)
Yellow fever (YF)
Yellow fever virus (YFV)
Zika virus (ZIKV)

## Acknowledgements

We would like to thank Dr Seth Irish for his continued support, expert knowledge and guidance throughout the entirety of the project. We would like to thank Siraj, Sintayheu and Getinet from the Centre for Public Health Laboratory, Jinka, Ethiopia and Johannis Negash from Public Health Emergency Management (PHEM) for their support, guidance and logistics during the field study. We would like to thank Cheryl Whitehorn for her support in morphological identification, and Yvonne Linton from Walter Reed Biosystematics Unit at the Smithsonian Institution for providing control mosquito samples. Finally, we would like to thank all those from South Omo Zone who agreed to participate in this study.

## Supporting Information Legends

Supplementary File S1: Arbovirus screening assays including PCR primer/probes sequences and cycling conditions.

Supplementary File S2: Household questionnaire: English version.

Supplementary File S3: STROBE checklist.

